# Probing enzyme-dependent pseudouridylation using direct RNA sequencing to assess neuronal epitranscriptome plasticity

**DOI:** 10.1101/2024.03.26.586895

**Authors:** Oleksandra Fanari, Sepideh Tavakoli, Yuchen Qiu, Amr Makhamreh, Keqing Nian, Stuart Akeson, Michele Meseonznik, Caroline A. McCormick, Dylan Bloch, Howard Gamper, Miten Jain, Ya-Ming Hou, Meni Wanunu, Sara H. Rouhanifard

## Abstract

Chemical modifications in mRNAs, such as pseudouridine (psi), can control gene expression. Yet, we know little about how they are regulated, especially in neurons. We applied nanopore direct RNA sequencing to investigate psi dynamics in SH-SY5Y cells in response to two perturbations that model a natural and unnatural cellular state: retinoic-acid-mediated differentiation (healthy) and exposure to the neurotoxicant, lead (unhealthy). We discovered that the expression of some psi writers change significantly in response to physiological conditions. We also found that globally, lead-treated cells have more psi sites but lower relative occupancy than untreated cells and differentiated cells. Interestingly, examples of highly plastic sites were accompanied by constant expression for psi writers, suggesting trans-regulation. Many positions were static throughout all three cellular states, suggestive of a “housekeeping” function. This study enables investigations into mechanisms that control psi modifications in neurons and its possible protective effects in response to cellular stress.

## Introduction

RNA modifications are enzyme-mediated chemical changes to the canonical structure of RNA nucleotides. Over 170 types of RNA modifications have been discovered in all classes of RNAs^1^ and play roles in diverse biological processes such as RNA metabolism^2,3^, translational control^4–6^, gene expression^3,7^, splicing^3,8^ RNA-protein interactions^7^, and immune response^9^. Of all uridines (U) in mammalian mRNA, 0.2% - 0.6% are pseudouridine (psi)^10,11^, a similar frequency as the abundance of m^6^A with respect to all adenosines^12,13^. Psi is an isomer of uridine^14^ in which a new hydrogen bond is available to base-pair primarily with adenosine but also other nucleobases when in an RNA duplex for a stabilizing effect^15, 3,16,17^. This duplex stabilization is thought to modulate cellular interactions with proteins and other biomolecules^3,8^.

Various next-generation sequencing methods have been utilized for psi mapping in mRNAs^6,10,18–21^; however, these methods all require chemical mediators (i.e., CMC labeling and bisulfite conversion) combined with reverse transcription to cDNA before amplification and sequencing. We and others have recently developed algorithms to classify psi sites from nanopore direct RNA sequencing (DRS)^22–25^. Our method, Mod-*p* ID^25^, compares the frequency of systematic basecalling errors at the modification site to an *in vitro* transcribed (IVT) unmodified transcriptome^26^. Mod-*p* ID accounts for the sequence context surrounding individual psi modifications and coverage at a given site to determine a statistical probability of a modification and provides a lower-limit occupancy value. A major benefit to this method is that it can provide a relative occupancy when the same position within the transcriptome is compared across different conditions. However, a caveat of this and other nanopore-based methods for psi-calling is that the methods alone are insufficient to validate psi modifications, necessitating exhaustive orthogonal validation approaches such as synthetic controls^27–29^ or biochemical assays^30^. Another suitable route for transcriptome-wide validation is to utilize knockdown/knockout of psi synthases (PUS) and measure changes in psi occupancies^8,21^ at the sites matching the motif for respective PUS.

In this work, we aim to understand how psi occupancies in mRNAs of neurons respond to changes in cellular state. Dysregulation of genes encoding PUS enzymes is associated with neuronal impairment.^31,32^ However, the plasticity of specific psi sites in neurons remains unknown. We used SH-SY5Y cells as a neuron-like system to model changes in cellular state and assess whether psi distribution and occupancy levels adjust in response to the environment. SH-SY5Y cells continuously express markers similar to immature catecholaminergic neurons while maintaining the ability to divide^33^. To understand the plasticity of psi, transcriptome-wide, we applied two different perturbations: one that models a healthy change in cellular state and one that models an unhealthy change in cellular state. The rationale for employing two distinct perturbations to investigate plasticity is that their mechanistic divergence would facilitate the identification of universally changing sites, stable sites, and those that may be condition dependent.

First, we exposed SH-SY5Y cells to retinoic acid (RA) to achieve differentiation, which models a healthy change in cellular state ^34^. Upon RA-mediated differentiation, SH-SY5Y cells become morphologically similar to primary neurons; their proliferation rate is decreased (similar to mature neurons), and the activity of enzymes specific to neuronal tissues is increased^35^. Second, we exposed them to lead (Pb^2+^), a systemic toxicant affecting virtually every organ system, but primarily the central nervous system^36^. Pb^2+^ is an environmental toxin that has been shown to adversely affect neuron functionality, especially during the developmental stages of the human brain.^37,38,39^

We performed DRS and Mod-*p* ID analysis to identify psi sites across conditions. To validate the identified sites as psi and assign them to specific psi synthases, we performed a siRNA knock-down of two key PUS enzymes, TRUB1 (motif GUUCN)^40^ and PUS7 (motif UNUAR)^41^. In addition to knockdown experiments, we performed rigorous validation of the query sites using five orthogonal methods: CeU-Seq^10^, BID-seq^21^, PRAISE^18^, CLAP^30^, and RBS-Seq^20^. These methods include RNA bisulfite sequencing, CMC-based sequencing, and biochemical CMC analysis by gel. Identifying a site as knocked down and through multiple orthogonal methods provides confidence that these sites are indeed psi.

We performed differential analysis, i.e., comparing psi occupancy at psi sites in untreated vs RA-differentiated cells, and untreated vs Pb^2+^-treated SH-SY5Y cells. Comparing three very different states can reveal the relative plasticity of psi sites at steady state (i.e., untreated SH-SY5Y cells) and in response to environmental cues for both differentiation and Pb^2+^ exposure. Such plasticity has been observed with m^6^A, for which modification levels increase significantly throughout brain development, which is suggested as a mechanism to achieve higher-order brain function^42^.

## Results

### Retinoic acid-induced differentiation of SH-SY5Y cells leads to a change in cell state

SH-SY5Y cells were differentiated into neuron-like cells by supplementing them with retinoic acid^43^, according to Kovalevich et al.^44^ (**Fig. 1a**). To confirm differentiation, we compared the cellular morphology between conditions and observed the characteristic elongation and branching of neurite-like processes from the differentiated cells (**Fig.1b**). Poly-A selected RNA was extracted from differentiated and untreated SH-SY5Y cells (n = 3 biological replicates for each group) and sequenced on a MinION device using R9 flow cells. We observed differential mRNA expression when comparing the two groups, supporting a change in cell state (**Fig 1d**). Importantly, we observed the expected differential mRNA expression of known differentiation markers^34,45,46^*. CRABP2*, *RARB*, *RGS2*, *RET*, and *DKK2* exhibited upregulation, and *ISOC1*, *MYC*, *SPRY2*, and *ASCL1* displayed decreased RNA expression in differentiated SH-SY5Y cells compared to the untreated counterparts (**Supplementary** Fig. 1). To assess the effects of retinoic-acid-mediated differentiation on psi machinery, we evaluated expression levels for 13 different PUS enzymes from untreated and differentiated libraries (**Fig. 1c**). While most PUS enzymes remained constant between conditions, we observed a significant difference in the RNA expression levels for *RPUSD3* and *PUS7L* in response to differentiation (*p* < 0.05). Rpusd3 is a pseudouridine synthase that is thought to act on mitochondrial RNA and ribosomal RNAs^47^. While the exact function of Pus7L is unknown, it may be regulated in response to physiological conditions as evidenced by tissue specificity of *PUS7L* in different brain tissues, as compared to the homolog *PUS7* which is ubiquitously expressed^48^.

**Figure 1.**
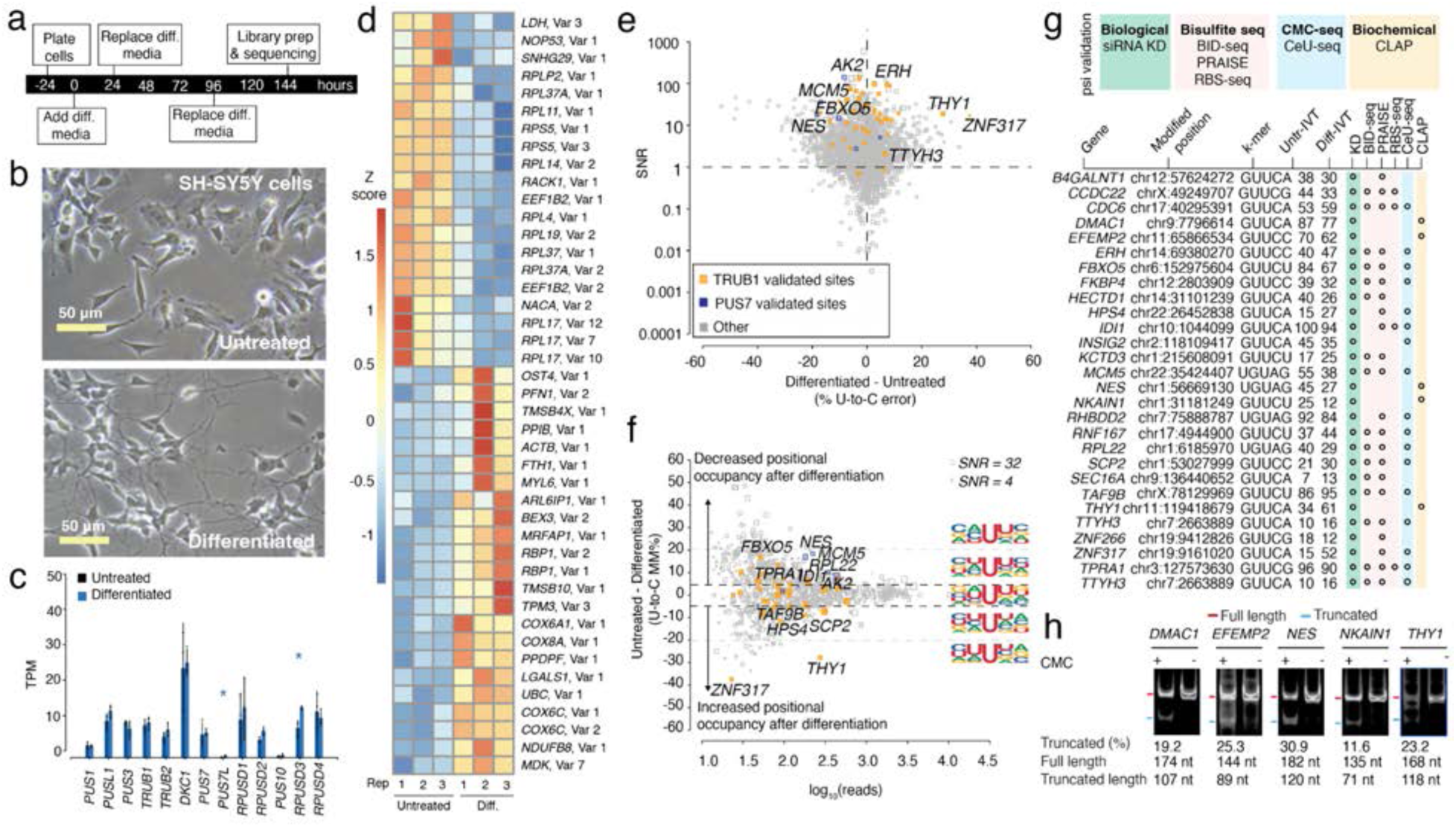
Effects of RA-mediated differentiation on mRNA psi modification and machinery in SH-SY5Y cells. a. Timeline illustrating the stages and duration of the RA treatment applied to SH-SY5Y cells. b. A representative photomicrograph of untreated and differentiated SH-SY5Y cells is shown. c. TPM of various PUS enzymes in untreated and differentiated SH-SY5Y cells determined by DRS. Individual colored bars represent each experimental condition, with error bars describing the standard error of the mean (SEM) across downsampled replicates. Individual replicates are shown as black dots. Statistics are performed by Student’s t-test, comparing each KD group to the scrambled control sample. * p < 0.05, ** p < 0.01, *** p < 0.001. d. We used Deseq2 to identify the transcripts with the highest fold change between the untreated and differentiated samples. Three biological replicates for each condition were used. The color scale shows a Z-score based on the relative fold-change. e. SNR vs. the difference in U-to-C error % between the untreated and differentiated samples. Orange dots represent uridine positions that are validated TRUB1 substrates, and blue dots represent uridine positions that are validated PUS7 substrates. f. Putative psi-positions determined by Mod-*p* ID are plotted according to the difference in U-to-C basecalling error in the untreated and differentiated samples against the reads for each position. Dotted line at the +5% and -5% marks indicate the cutoff for a position to be changed in response to perturbation. Inset shows the sequencing logo for positions within the TRUB1 motif, gray points above the threshold line, and total points above the threshold line. g. Annotation of genes containing a psi modification validated by PUS7 or TRUB1 KD (Figure 2) and orthogonal methods. h. CLAP gel result of psi incorporation in mRNAs for the sites confirmed by Nanopore DRS and KD data, but not by any other orthogonal method. See Supplementary Fig. 2 for original gel source data.

### Transcriptome-wide mapping of psi-modifications before and after differentiation

Psi positions were identified in differentiated and untreated SH-SY5Y cells by enriching our paired *in vitro* transcribed (IVT) dataset with additional human IVT data^26^ (see **Methods**), allowing us to achieve sufficient coverage for an additional 878 putative psi sites. We then used Mod-*p* ID^25^ to identify putative psi sites based on significant differences in U-to-C basecalling error (*p* < 0.001) in the untreated and differentiated libraries compared to the IVT control library.

Sites with > 40% U-to-C error (i.e., hypermodification type I^25^) may be considered “high-occupancy” and have the potential for stronger phenotypes. These sites were assessed across samples (**Supplementary Table 1**). It should be noted that the percentage here is only a qualitative metric of occupancy. From the untreated sample, 131 hypermodified sites were detected, and 132 sites were detected from the differentiated library. (**Supplementary** Fig. 2a, b). Of these sites, 83 were hypermodified in both conditions (**Supplementary** Fig. 2d). Most of these sites fall within mRNAs that encode proteins of various functions. Notably, for the sites that are unique to differentiated samples, *CAMKV*, *KIF2A*, *DBNL*, and *CRK*L are involved in neuronal and synaptic function.

The occupancy levels of sites common to both libraries were compared to identify differences in psi profiles due to the differentiation process beyond just the hypermodified sites. First, we selected positions present in both the untreated and differentiated libraries and defined a signal-to-noise ratio (SNR) for each site (see **Methods**). This filtering step was introduced to guarantee a strong DRS signal level as compared to the IVT while considering coverage and size of error signatures (**Fig 1e**). We selected sites with SNR ≥ 1 and read coverage of ≥ 10 reads in both libraries (**Fig. 1f**). We observed that 1,786 sites exceeded this threshold, indicating that the signal-to-noise ratio was sufficient for analysis (**Supplementary Table 2**).

Next, we performed a differential analysis comparing the relative occupancy of identified sites in the untreated and differentiated libraries. We found a mean difference in U-to-C mismatch of 2.46% with a 95% CI = [2.02, 2.91], indicating a statistically significant change in modification occupancy between the differentiated and untreated SH-SY5Y cells. This result suggests that the relative modification occupancy is lower in differentiated SH-SY5Y cells compared to the untreated cells for conserved targets. We focused on sites with a differential U-to-C mismatch level outside the CI. To make our differential analysis more conservative, we defined sites as “changed” if samples had observed a ≥ 5% difference in U-to-C error in both directions (**Fig. 1f**).

To explore whether a specific sequence motif is changed, we exported the sequencing logo for positions categorized into three groups: positions with higher relative occupancy following differentiation, positions that remain unchanged, and positions with lower relative occupancy following differentiation (**Fig. 1f**). For the positions in which the positional occupancy decreased after differentiation, the +1-nucleotide neighboring the psi position was frequently uridine while the unchanged and higher occupancy sites equally represented all nucleotides in this position.

Among the sites with significant differences between samples (i.e., sites that fell outside of the CI), 26 positions were assigned to a specific PUS enzyme by siRNA mediated knockdown (KD) of *PUS7* and *TRUB1* (**Supplementary Table 3**). 86% of these positions were additionally validated as psi using our siRNA KD with at least 1 out of these four orthogonal methods: BID-Seq^21^, PRAISE^18^, RBS-seq^20^, and CeU-Seq^10^. We found that five positions (*DMAC1* chr9:7796614, *EFEMP2* chr11:65866534, *NES* chr1:56669130, *NKAIN1* chr1:31181249, *THY1* chr11:119418679) were validated by our knockdowns but have not been discovered by these four orthogonal methods (**Fig. 1g**) so we proceeded to validate these sites with CLAP (CMC-RT and ligation assisted PCR analysis of psi modification)^30^. CLAP confirmed all these sites identified that were only by Mod-*p* ID as psi serving as orthogonal validation (**Fig. 1g-h, Supplementary** Fig. 3).

The most substantial changes were for *ZNF317* (chr19:9161020), which increased from a 15% U-to-C error in the untreated sample to 52% in the differentiated sample. The next most substantial change was for *THY1* (chr11:119418679), which increased from a 34% U-to-C error in the untreated sample to a 61% U-to-C error in the differentiated sample. The modified position for *ZNF317* is within the CDS and the encoded protein is part of the zinc finger protein family and is involved in transcriptional regulation and cellular adaptation to changes^49^. The modified position for *THY1* is in the 3’UTR, and the encoded protein is a cell surface glycoprotein involved in cell adhesion processes and modulates neurite outgrowth^50^. An example of a reduced psi occupancy upon differentiation (25% to 12%) is found for *NKAIN1,* which encodes a protein involved in sodium-potassium transport in the brain^51^.

Next, we explored differences in mRNA expression for transcripts that harbor a psi position. Interestingly, the mRNA levels remained unchanged, with differences in psi occupancy. (**Supplementary** Fig. 4a) We found that only *IDI1* (chr10:1044099; 15.3 TPM in the untreated sample and 58.9 TPM in the differentiated sample) showed a significant difference in mRNA expression between the two conditions (*p* < 0.001). Other methods have also reported that this site is a substrate for Trub1^18^. *IDI1* encodes an enzyme that is involved in the synthesis of cholesterol metabolites which are essential for the proper differentiation and function of neurons. Next, we examined the protein expression levels in cellular compartments for the two dominant PUS enzymes for humans, Pus7 and Trub1, using immunofluorescence in untreated and differentiated SH-SY5Y cells. Our analysis revealed no significant differences in the subcellular distribution of these two PUS enzymes (**Supplementary** Fig. 5).

### Absolute quantification of psi occupancy in response to differentiation

While our differential analysis highlights sites with significant relative changes in occupancy, we were interested in assigning absolute occupancy to some of these positions. Sequence-specific synthetic controls were generated to match the mRNA sequences bearing psi in the undifferentiated and differentiated samples. One library with uridine in the modification site and one with psi in the modification site was generated and sequenced using DRS. Using these sequence-specific synthetic controls, we trained 10 supervised machine-learning models (ML) to quantify the occupancy of 10 psi-sites (see **Supplementary Table 4**) in the native RA-differentiated and untreated transcripts (see **Methods**) using ModQuant^29^. Each ML model had a classification accuracy of > 90%. *PSMB2* (chr1:35603333) is an example of a site with similar occupancies in the untreated (62% ML-predicted level, 61% U-to-C mismatch) and differentiated library (61% ML-predicted level, 63% U-to-C mismatch). An opposite example is *PRPSAP1* (chr17:76311411), which changes both in the untreated (67% ML-predicted level, 47% U-to-C mismatch) and differentiated sample (62% ML-predicted level, 33% U-to-C mismatch). This highlights the importance of generating synthetic controls for absolute quantifications.

When applied to the DRS libraries, the lowest level of occupancy predicted by the ML models is 16.9% (ML model for *MRPS14*). These stoichiometry levels are expected as the sites we tested with synthetic standards have at least 20% U-to-C mismatch in the native datasets. However, this approach can reach single-read classification accuracy of 95%, as shown by the model error calculated on each synthetic standard (**Supplementary** Fig. 6).

### SH-SY5Y cell state changes in response to lead (Pb^2+^) exposure

Next, we introduced the neurotoxicant Pb^2+^ to SH-SY5Y cells for six days as an alternative, “unhealthy” change in cellular state to contrast with differentiation as a “healthy” change (**Fig. 2a**). Previous studies have shown that 5 μM is close to the Pb^2+^ levels in human blood that can cause encephalopathy in children^53^; however, we chose to use a higher Pb^2+^ concentration (50 μM) because the *in vitro* tolerance for cytotoxicity is higher than *in vivo*^53^ while maintaining cell viability and typical cellular morphology following exposure (**Fig. 2b**). Poly-A selected RNA was extracted from untreated and Pb^2+^-exposed SH-SY5Y cells (n = 2 biological replicates) and sequenced on a MinION device using R9 flow cells. Global gene expression profiling of untreated and Pb^2+^-exposed libraries support a change in cellular state (**Fig. 2d**). To assess the effects of Pb^2+^ exposure on psi machinery, we evaluated mRNA expression levels for 13 different psi-synthases from Pb^2+^ treated and untreated SH-SY5Y cell libraries, and we found no significant differences in expression levels (**Fig. 2c**).

**Figure 2.**
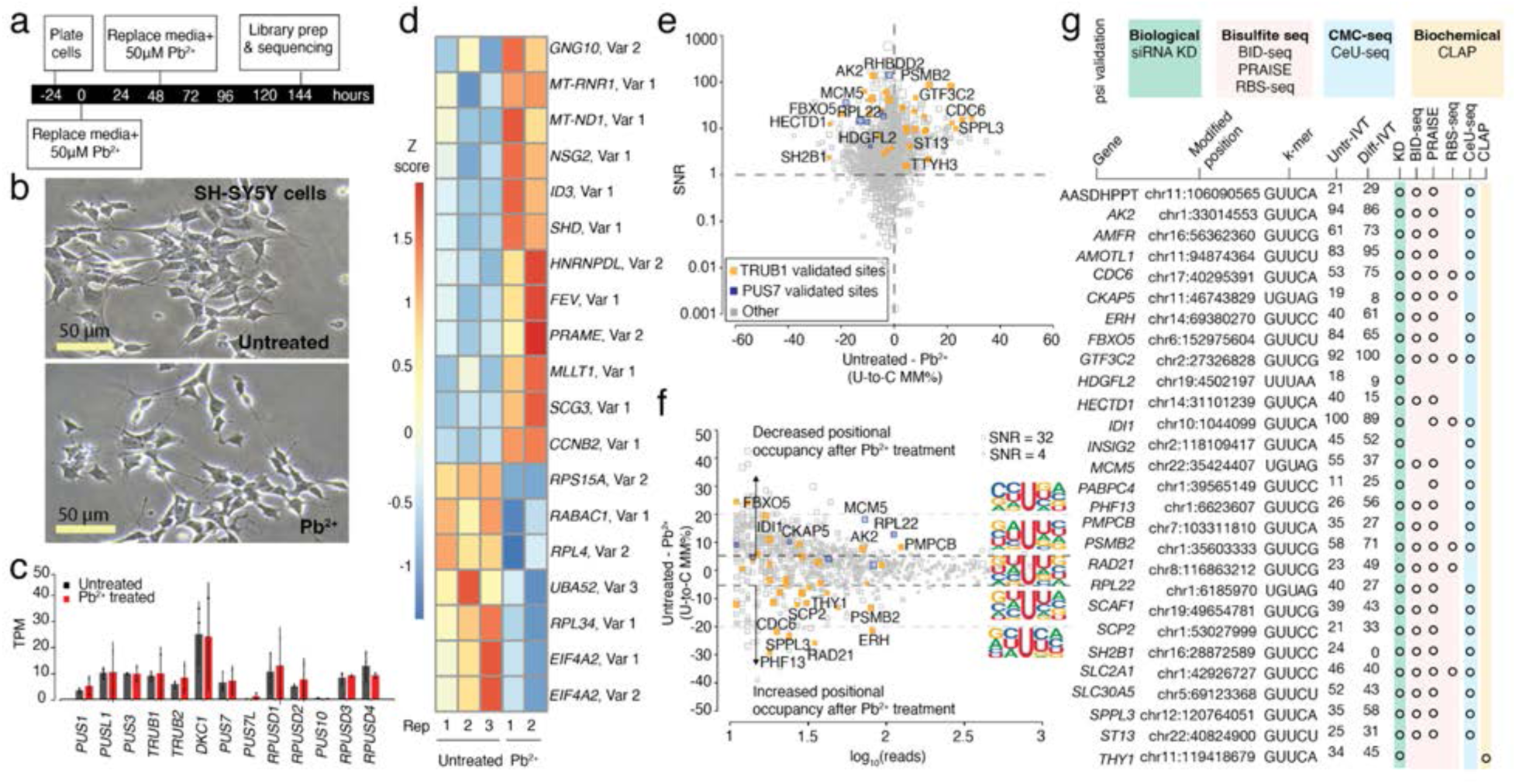
Effects of Pb^2+^ treatment on mRNA psi modification and machinery in SH-SY5Y cells. a. Timeline illustrating the stages and duration of the Pb^2+^ treatment applied to SH-SY5Y cells. b. A representative photomicrograph of untreated and Pb^2+^ treated SH-SY5Y cells is shown. c. DRS determined the TPM of various PUS enzymes in untreated and Pb^2+^-treated SH-SY5Y cells. Individual colored bars represent each experimental condition, with error bars describing the standard error of the mean (SEM) across downsampled replicates. Individual replicates are shown as black dots. Statistics are performed by Student’s t-test, comparing each KD group to the scrambled control sample. * p < 0.05, ** p < 0.01, *** p < 0.001. d. We used Deseq2 to identify the transcripts with the highest fold change between the untreated and Pb^2+^ treated samples. Three biological replicates of each condition were used. Color scale shows a Z-score based on the relative fold-change. e. SNR vs. the difference in U-to-C error % between untreated and Pb^2+^ treated samples. Orange dots represent uridine positions that are validated TRUB1 substrates, and blue dots represent uridine positions that are validated PUS7 substrates. f. Putative psi-positions determined by Mod-*p* ID are plotted according to the difference in U-to-C basecalling error in the untreated and Pb^2+^ treated samples against the reads for each position. A dotted line at the +5% and -5% marks indicate the cutoff for a position to be changed in response to perturbation. The inlet shows the sequencing logo for positions within the TRUB1 motif, gray points above the threshold line, and total points above the threshold line. g. Annotation of genes containing a psi modification that changed in response to perturbation and validated by PUS7 or TRUB1 KD (Figure 2) and orthogonal methods.

### Transcriptome-wide mapping of psi-modifications following Pb^2+^ exposure

To explore changes in psi modification between untreated and Pb^2+^ treated cells, we generated a paired, unmodified transcriptome for untreated SH-SY5Y cells to identify putative psi positions. We enriched the paired IVT (see **Methods**), recovering 333 sites that would have been discarded because of insufficient coverage in the paired IVT^26^. We then applied Mod-*p* ID^25^ to identify putative psi sites based on significant differences in U-to-C basecalling error (p < 0.001 in at least two biological replicates, see **Methods**). First, the frequency of sites with > 40% U-to-C error (i.e., hypermodification type I^54^) was assessed. We detected 74 hypermodified sites from the Pb^2+^-treated library, of which 48 were shared with the untreated library (**Supplementary** Fig. 2c, d). We selected positions found in both the untreated and Pb^2+^ treated libraries and calculated the SNR for each site as described above (**Fig. 2e**), obtaining 946 sites that match our filtration criteria. From these sites, we found a mean difference in U-to-C mismatch error of 2.72% and a 95% CI = [2.14,3.29], indicating a statistically significant change in modification occupancy between the Pb^2+^-treated and untreated SH-SY5Y cells. This result indicates that the global modification occupancy is lower in Pb^2+^-treated cells compared to the untreated cells for conserved mRNA targets. Many sites unique to the Pb^2+^ treated sample fall within mRNAs encoding proteins involved in oxidative stress response, including *MRPS14*, *DHCR7*, *RTN4*, *SIGMAR1*, and *SAE1*.

Sites were defined as changed in response to Pb^2+^ treatment using the same CI-based criteria applied to RA-differentiated SH-SY5Y cells explained above (**Fig. 2f, Supplementary Table 5**). We exported the sequencing logo for positions categorized into three groups: positions with higher relative occupancy following Pb^2+^ treatment, positions that remained unchanged, and positions with lower relative occupancy following Pb^2+^ treatment (**Fig. 2f**). We found that psi sites with increased positional occupancy following Pb^2+^ treatment tend to be flanked by two uridines, while those with decreased positional occupancy following Pb^2+^ treatment have a uridine in the n+1 position.

Among the sites with significant differences between samples, we found 28 positions that were assigned to a specific PUS enzyme using our KD experiments (**Supplementary Table 7**). In addition to our siRNA KD validation, we confirmed these sites using the same orthogonal methods that were used for differentiated cells (**Fig. 2g**). Among these sites, 26 (93%) were validated with at least one additional orthogonal method. Two positions, *HDGFL2* (chr19:4502197) and *THY1* (chr11:119418679) transcripts, were not validated by previously annotated datasets so we validated using the CLAP method. The site within *HDGFL2* was not possible to test with this method because it is only 10 nucleotides away from the end of the 3′ UTR, leaving insufficient space to design CLAP primers. We were able to apply the method to *THY1*, which was confirmed by CLAP as a psi-modified site (**Fig. 1h, Supplementary** Fig. 3). In the comparison of psi sites in untreated and Pb^2+^ treated samples, the biggest change was for *PHF13* (chr1:6623607), which increased from 26% U-to-C error in the untreated library to 56% U-to-C error following Pb^2+^ treatment. Phf13 modulates chromatin structure and DNA damage response.

We calculated TPMs for each Trub1 and Pus7 mRNA substrate with differences in psi levels to explore differences in mRNA expression for transcripts that harbor a psi. We found that only *ERH* (chr14:69380270; 158 TPM in the untreated sample and 207 TPM in the Pb^2+^ treated sample) showed a significant difference in mRNA expression between the two conditions (*p* < 0.05; **Supplementary** Fig. 4b). We then examined the protein expression levels in cellular compartments for the two dominant PUS enzymes for humans, Pus7 and Trub1, using immunofluorescence in untreated and Pb^2+^ treated SH-SY5Y cells and found no significant differences in the subcellular distribution of these two PUS enzymes (**Supplementary** Fig. 7).

### Absolute quantification of psi occupancy in response to Pb^2+^ exposure

Following the same approach described for RA-differentiation stoichiometry assessment, we used ModQuant^29^ and trained 9 supervised ML models to quantify the occupancy of 9 psi-sites (see **Supplementary Table 4**) in the transcripts from the Pb^2+^-treated and untreated libraries (see **Methods**). As in the differentiated library, *PSMB2* (chr1:35603333) is an example of a site with similar occupancies in both untreated (62% ML-predicted level, 61% U-to-C mismatch) and lead-treated library (78% ML-predicted level, 74% U-to-C mismatch). Alternatively, U-to-C mismatch and ML-predicted occupancy were different for *SLC2A1* (chr1:42926727) in both untreated (67% ML-predicted level, 46% U-to-C mismatch) and lead-exposed samples (74% ML-predicted level, 40% U-to-C mismatch).

### Plasticity of pseudouridylation of mRNAs in response to changes in the cellular state

We then sought to compare our 3 cellular states for the same cell type to see which psi positions are static and which are plastic (i.e., cell state dependent psi regulation). We compared the relative occupancy for psi-modified positions across perturbations. We selected orthogonally-validated positions that met the following criteria to evaluate the changes to individual positions between the three conditions: 1. Detected by Mod-*p* ID as a psi position with *p*-value < 0.001 in at least one condition; 2. U-to-C mismatch was >40% in at least one of the conditions; 3. Number of reads in all conditions was >10 in DRS and IVT libraries. We rank-ordered these 36 positions based on the standard deviation (SD) over the three conditions (**Fig. 3a**). Interestingly, 27 out of these 36 positions fall within a TRUB1 motif, and 5 fall within a PUS7 motif. The most similar position between the three conditions is *INTS1* (chr7:1476655), with a SD of 0 and >90% U-to-C error for each condition. *INTS1* is a component of the integrator complex which has been linked to developmental delays and is involved in in RNA processing and transcriptional regulation^55–57^. Positions with an SD of >5 are considered to have relatively high. Among these sites, we identify 10 positions with a SD > five, which we consider the most static psi positions. The least similar position is on *THY1* (chr11:119418679), with a SD of 19.6 and U-to-C errors of 33.5%, 61.3%, 45.2%, on untreated, differentiated, and Pb^2+^-treated libraries respectively. *THY1* encodes a cell surface glycoprotein involved in cell adhesion processes and modulates neurite outgrowth^58^.

**Figure 3.**
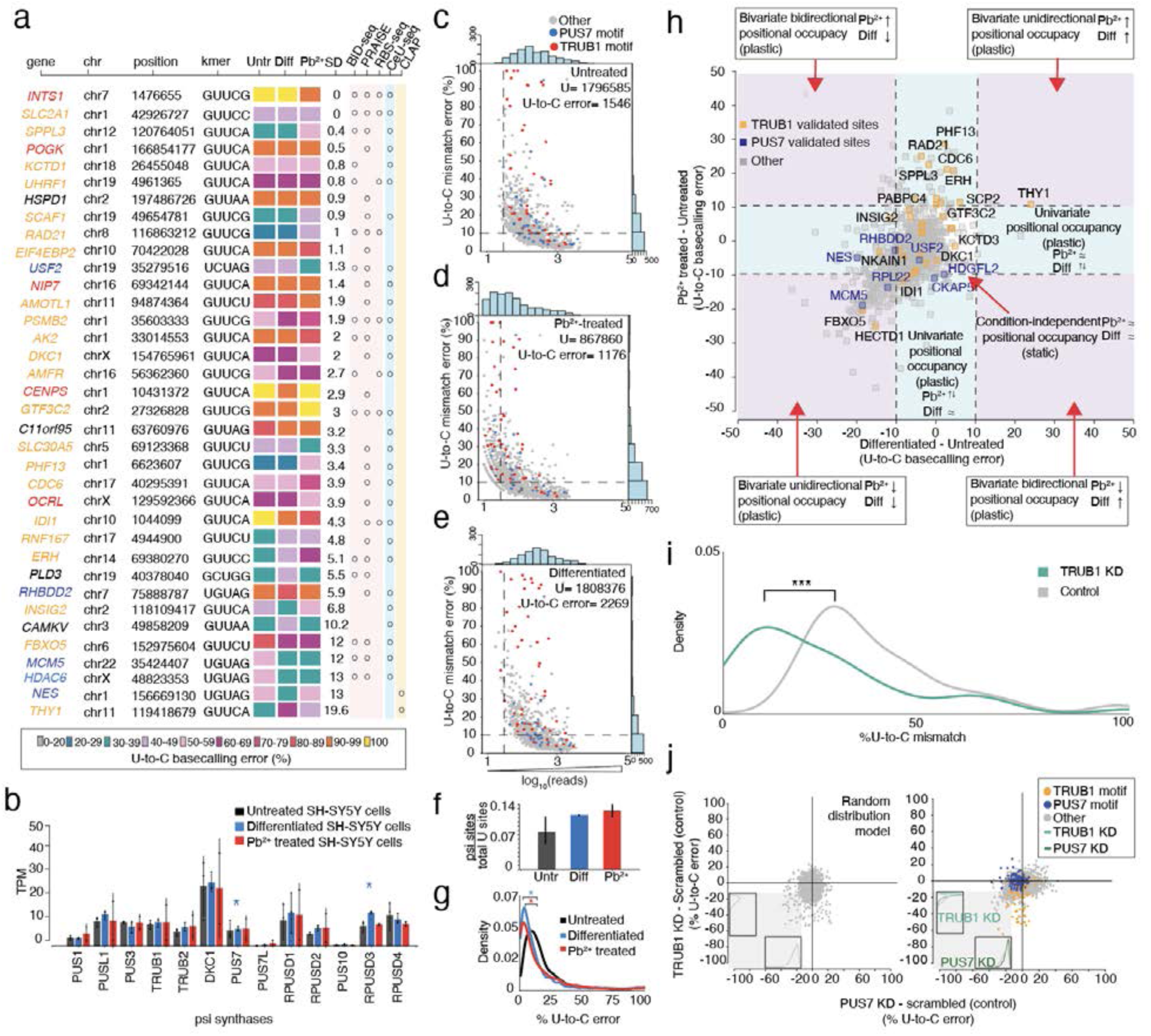
Psi analysis across three cellular states enables the classification of plastic and static sites of modification, transcriptome wide. a. Heatmap of sites with at least 40% U-to-C basecalling error in one of three conditions. Colors indicate the percentage of U-to-C basecalling errors. For each position the standard deviation (SD) is calculated on all three conditions and used to rank the sites in order, with the most similar at the top and the least similar at the bottom. KD validated substrates for TRUB1 are shown in orange, and TRUB1 motifs that have not yet been validated are shown in red. KD validated substrates for PUS7 are shown in dark blue, and PUS7 motifs that have not been validated are shown in light blue. Orthogonal validation with non-nanopore methods is performed for all the sites. b. DRS determined the TPM of various PUS enzymes in untreated, differentiated, and Pb^2+^-treated SH-SY5Y cells. Individual colored bars represent each experimental condition, with error bars describing the standard error of the mean (SEM) across downsampled replicates. Individual replicates are shown as black dots. Statistics are performed by Student’s t-test, comparing each KD group to the scrambled control sample. * p < 0.05, ** p < 0.01, *** p < 0.001. c. Untreated SH-SY5Y number of total DRS reads are plotted against the U-to-C mismatch error of putative Ψ sites detected by Mod-*p* ID (p-value < 0.001 in at least two biological replicates). Blue dots represent uridine positions within a PUS7 motif, while red dots represent uridine positions within a TRUB1 motif. All the other motifs are shown in grey. Dashed lines represent the criteria used for defining a position as a psi site (number of reads>10; U-to-C error>10). d. Pb^2+^ treated SH-SY5Y number of total DRS reads are plotted against the U-to-C mismatch error of putative Ψ sites detected by Mod-*p* ID (p-value < 0.001 in at least two biological replicates). e. Differentiated SH-SY5Y number of total DRS reads are plotted against the U-to-C mismatch error of putative Ψ sites detected by Mod-*p* ID (p-value < 0.001 in at least two biological replicates). f. Colored bars shows the number of significant pseudouridine sites, labeled as U-to-C error in panels d,e,f, normalized by the total number of reads in each of the three conditions. Error bars show the standard error of the mean across biological replicates. g. Distribution of the occupancy levels calculated on untreated, differentiated, and lead-treated libraries. Statistical significance is calculated using 95% CI, * p < 0.05. h. Putative psi-positions determined by Mod-*p* ID are plotted according to the difference in U-to-C basecalling error in the untreated and Pb^2+^ treated samples against the difference in U-to-C basecalling error between the differentiated and untreated for each position. A dotted line at the +10% and -10% marks indicate the cutoff for a position to be changed in response to perturbation. Condition-independent positional occupancy (static) sites are shown in the center square. Single condition-dependent positional occupancy sites are shown in the blue stripes. Double condition-dependent positional occupancy sites are shown in the purple areas. i. Distribution of the occupancy levels calculated on PUS7 validated sites before (grey) and after the TRUB1 KD (green). The occupancy levels in the KD library are decreased compared to the control condition without KD. Statistics are performed by Student’s t-test, comparing each KD group to the scrambled control sample. * p < 0.05, ** p < 0.01, *** p < 0.001. j. Co-operation effect of TRUB1 and PUS7 KD on SH-SY5Y cells. (Left) Random distribution model of points constrained by the data parameters. Distributions are calculated for points in the lower left quadrant as a function of each simulated knockdown. (Right) distribution of TRUB1 KD plotted against PUS7 knockdown. Distributions are calculated for points in the lower left quadrant as a function of each knockdown.

The abundance of psi synthase may account for some of the differences in psi percentage within that condition. Hence, we compared mRNA levels of psi synthases across all three conditions and found significant differences in PUS7 and RPUSD3 expression (**Fig. 3b**). We also hypothesized that for the sites with higher plasticity, mRNA expression of the target may explain the discrepancy (**Supplementary** Fig. 4c). However, we did not observe an association between mRNA expression levels and the percentage of U-to-C error for targets with >10 SD.

To assess which cell condition had the highest number of sites bearing psi (irrespective of occupancy), we calculated the number of significant psi sites (*p*-value < 0.001 in at least two biological replicates) with sufficient coverage (>10 reads in DRS and IVT libraries) detected by Mod-*p* ID across cell conditions and normalized it by the total number of reads in each of the three samples (see **Methods**). We found that the Pb^2+^-exposed cells have the higher normalized percentage of psi sites (0.14%), followed by the differentiated (0.13%) and untreated conditions (0.09%; **Fig. 3c-f**). Interestingly, for the sites shared between the three libraries, we find a statistically significant decrease in occupancy levels (*p*<0.05; **Fig. 3g**) in both treated libraries (differentiated and Pb^2+^) compared to the untreated one.

Finally, we divided the positions into different zones based on the relationship between the conditions: 1. Univariate positional occupancy whereby the relative occupancy of a given site changes for either differentiation or Pb^2+^ exposure, demonstrating the plasticity of a given site; 2. Bivariate positional occupancy whereby the relative occupancy changes for both differentiation and Pb^2+^ exposure, demonstrating plasticity; 3. Condition-independent positional occupancy, whereby the occupancy does not change in response to the different conditions (these modifications are present and stable between different perturbations, i.e., static; **Figure 3g**).

We found that 73% of sites are static (condition 3) and 27% are plastic (condition 1 and/or 2; **Supplementary Table 7**). Most of the Trub1 targets fell within condition 3 (static). Notably, we found two TRUB1 KD validated sites in the univariate group affected by differentiation and ten TRUB1 KD validated psi sites in the univariate Pb^2+^-dependent group. These include sites within *PHF13*, *CDC6* and *RAD21,* which encode proteins that are linked to DNA repair mechanisms and cellular response to toxicants^59^. These are examples of potentially interesting sites for future analysis as mediators of RNA binding proteins or translational control.

Interestingly, we observed a cluster of targets (541 sites) that were decreased for both Pb^2+^ exposure and differentiation which we classified as group 2 plasticity. This cluster is interesting because these targets are higher occupancy in untreated cells and decreased in both perturbations. We compared this distribution to a random distribution and found that the two distributions were significantly different (Mann–Whitney–Wilcoxon test, p < 0.001), meaning that the concomitant decrease in both perturbations is inconsistent with a random effect. Among these plastic sites with a bivariate, unidirectional decrease in occupancy, we find sites within *MCM5* and *FBXO5*, involved in cellular stress response; *HECTD1,* which is known to modulate retinoic acid signaling^60^ and *RPL22*, a ribosomal protein linked to common cancer-associated mutations and nucleolar stress response^61,62^.

### Modeling co-regulation of TRUB1 and PUS7 on mRNA substrates

We observed that a group of psi-sites that were validated as PUS7 substrates showed decreased occupancy levels in the TRUB1KD library (i.e., the TRUB1 KD determines a lower psi occupancy not only on TRUB1 motifs but also within PUS7 motifs; **Supplementary** Fig. 8d). To test if TRUB1 KD causes an occupancy reduction not just in TRUB1, but also in PUS7 sites, we plotted the distribution of the %U-to-C mismatch of validated PUS7 sites pre- and post-TRUB1 KD. We found a statistically significant shift of PUS7 sites (UNUAR motif) towards lower occupancy levels in the TRUB1 KD library (*p*=0.0003, Student’s t-test; **Fig. 3i**). This shift towards lower occupancy levels is unexpected, as it occurs in TRUB1 KD samples. In fact, we would expect this to happen upon the KD of PUS7 as this is the writer acting on UNUAR motifs. This suggests that the TRUB1 and PUS7 enzymes have a co-regulatory mechanism.

To explore this observation further, we first assessed the changes in PUS mRNA levels upon TRUB1 and PUS7 KD and observed a significant reduction in *TRUB1* mRNA upon PUS7 KD (Student’s t-test, p-value < 0.05) and a reduction in *PUS7* upon TRUB1 KD compared to the scrambled control (**Supplementary** Fig. 8c).To test the hypothesis that TRUB1 KD has a global effect on psi levels within PUS7 motifs, we compared the differences in positional occupancy for the TRUB1 KD with the PUS7 KD to a random distribution model of positions that were constrained on the parameters of the observed sequencing data (**Fig. 3j**). We found that the data are inconsistent with a random model (Mann–Whitney–Wilcoxon test, p < 0.001) in the third quadrant, a region where the position shows a decreased positional occupancy for both TRUB1 KD and PUS7 KD. We repeated the same analysis on two additional random models, both statistically different from the real dataset (Mann–Whitney–Wilcoxon test, p < 0.01; **Supplementary** Fig. 9a). To support a potential coregulation of TRUB1 and PUS7 that we observed on SH-SY5Y cells, we applied the same analysis and filtering steps to a HeLa cell dataset on which we performed a siRNA-based knockdown of the same enzymes TRUB1 and PUS7 (**Supplementary** Fig. 9b). As described before, we randomized the distribution of the real data constraining it to the parameters of the real sequencing data. As for SH-SY5Y cells, we found that HeLa KD data are not consistent with a random model (Mann-Whitney-Wilcoxon test, p < 0.001) in the decreased-occupancy region. These data support a possible coregulation of Trub1 and Pus7.

## Discussion

This study assessed the plasticity of psi machinery and modifications in SH-SY5Y cells in response to perturbation of cellular state. We found that, in response to differentiation, the mRNA levels of psi writers *PUS7L* and *RPUSD3* were increased. *PUS7L* has been shown to have cell type-specific expression in the brain while *PUS7* has relatively stable expression across cell types. This observed change is the first experimental evidence that *PUS7L* is regulated in response to physiological conditions because the same neuronal cell type is compared in 2 different conditions.

Our comparative analysis of the positional occupancy in multiple cellular conditions (untreated, differentiated, and Pb^2+^ treated) revealed various types of site responses: we found numerous positions with high variable occupancy across conditions, indicating plasticity at those sites. Interestingly, the most static psi positions were frequently targets of TRUB1, which suggests a high degree of conservation both in the positions detected and the occupancy. We also found that 3 out of 4 PUS7 sites were in the plastic group, which may indicate that PUS7 deposited sites are less conserved across conditions although this may be due to a low number of reads. Among these, *NES* (chr1:156669130) encodes the neuronal marker and cytoskeletal protein Nestin, which plays a role in differentiation and self-renewal. This *NES* psi site has decreased psi levels following differentiation, while it does not change upon Pb^2+^ treatment. Noteworthy, mRNA levels for *NES* are unchanged across all conditions (**Supplementary** Fig. 4c). In contrast, *THY1* (chr11:119418679) exhibits high upregulation of psi occupancy for both Pb^2+^ treatment and differentiation and encodes for a glycoprotein involved in neuronal processes. Possible functions for sites highly sensitive to environmental factors include translational control in response to cellular stress and maintenance of cellular fitness. Future studies may test cellular fitness in the presence of these stressors in combination with TRUB1/PUS7 knockdown to determine mechanism.

Among the most static positions, some psi sites are found on genes linked to neuronal functions: *SLC2A1* is associated with GLUT1 deficiency syndrome, a neurological disorder characterized by seizures, developmental delay, and movement disorders^63,64^. Dysregulation of *UHRF1* has been implicated in various cancers and neurodevelopmental disorders^61,62^. *EIF4EBP2* regulates translation initiation by binding to eukaryotic translation initiation factor 4E (eIF4E) and inhibiting its interaction with the mRNA cap structure. It plays a role in synaptic plasticity and memory formation and in neurodevelopmental disorders such as autism spectrum disorders^65–67^. Other transcripts, although not relevant to neuronal functions, contain psi static sites. The biological function of these genes is general and related to basic functions and cellular mechanisms. Future studies may test the hypothesis that static sites serve a “housekeeping” function, maintaining homeostasis.

Finally, a global pseudouridylation level assessment of each cell condition (untreated, differentiated, and Pb^2+^ treated) showed that the Pb^2+^ libraries had a higher normalized percentage of psi sites compared to the untreated (**Fig. 3f**). Interestingly, despite the higher frequency levels, we find a statistically significant decrease of the occupancy levels in differentiated and lead-treated libraries compared to the untreated libraries (p<0.05; **Fig. 3g**) despite the higher frequency levels (**Fig. 3f**). We hypothesize that the broader installation of the psi-sites across the transcriptome is used by the cell as a mechanism of protection/response to the treatment and that the enzyme responsible for installing the modification may not work as efficiently as in the untreated cells because of enzymatic alterations (i.e., saturation); however, further studies will be required to assess the hypothesis.

We validated psi sites by siRNA knockdown of the predominant psi synthases acting on human mRNAs, TRUB1 and PUS7. Interestingly, we found that TRUB1 and PUS7 KDs affect other psi writers. This effect is stronger in PUS7 KD cells, as the other six PUS enzymes (*RPUSD1*, *TRUB1*, *DKC1*, *PUSL1*, *RPUSD4*, *RPUSD3*) have a significant reduction in their mRNA levels, while TRUB1 KD only affects the RPUSD1 enzyme. This finding is suggestive that the Pus7 protein may act as a transcription factor for the other psi synthase enzymes. Interestingly, in TRUB1 KD libraries, certain positions displaying TRUB1 motifs with high U-to-C basecalling errors did not exhibit changes following TRUB1 KD, and most of these positions have been validated through chemical-based methods. It is possible that, with only a partial knockdown, there were sufficient levels of enzyme available to modify the site. Alternatively, other PUS enzymes could compensate for the decrease in Trub1 enzyme.

We also found evidence of co-regulation for PUS7 and TRUB1. This was a surprising finding, and we analyzed the KD datasets published by Dai et al.^21^ and found 21 psi sites to have reduced psi modification levels under the depletion of both TRUB1 and PUS7 enzymes, supporting our finding.

We used DRS-independent methods to confirm the knockdown-validated sites found in the differentiated and Pb^2+^-exposed libraries. We found that *DMAC1*, *EFEMP2*, *NES*, *NKAIN1*, *THY1, and HDGFL2* transcripts carried a psi site that had not been discovered by other orthogonal methods in the differentiated sample. We found four of these transcripts (*TPM3*, *PIR*, *HDGFL2*, and *NES*) in the Pb^2+^-exposed library. The presence of these unconfirmed sites could be because the SH-SY5Y cell line has not been previously analyzed by any other groups. This is further suggested by our CLAP results, which validated five new sites that we uncovered to be psi.

This study was the first to determine the plasticity of psi modifications across cellular states. Future analysis will determine whether static sites play critical roles in the cell’s biological function and whether the plastic sites are responses to external cues for fine-tuning gene expression.

## STAR Methods

### Experimental Model and Subject Details Cell culture

Human neuroblastoma SH-SY5Y cells were cultured in EMEM/F12 (EMEM from Quality Biological Inc and Cytiva HyClone Ham’s Nutrient Mixture F12 Media) supplemented with 10% Fetal Bovine Serum (FisherScientific, FB12999102). For untreated SH-SY5Y cells, the culture remained in this medium for seven days at 37C and 5% CO_2_, refreshed every three days. For differentiated SH-SY5Y cells, after 24h, the media changed to differentiation media, which is Neurobasal media (Gibco Neurobasal-A Medium, minus phenol red) supplemented with 10uM all-trans-retinoic acid (Fisher, AC207341000), 1X B27(Fisher, A3582801), and 1X Glutamax (Fisher, 35-050-061). The differentiation media was renewed every other day. For lead exposure SH-SY5Y cells, after 24 hours, the culture media was removed, and a 50 μM Pb^2+^-supplemented (Lead (II) acetate trihydrate, Sigma) untreated media was added to the cells. The media was replaced every three days.

### Immunofluorescence (IF)

For fixing SH-SY5Y cells, half of the culture media was removed, and an equal volume of 4% formaldehyde (Fisher, F79500) in PBS was added to each well for the final of 2% formaldehyde. After 2 min incubation at room temperature, the solution was aspirated, replaced by 4% formaldehyde, and incubated for 10 mins. The cells were then washed with PBS and permeabilized by incubating in PBS-Triton (0.1%) for 10 min. The cultures were blocked by incubation in 2% bovine serum albumin (BSA) in BS-Triton (0.1%) for one hour, followed by three times washed with PBS-Tween 20 (0.1%). The cells were then incubated with 1ug/ul primary antibody (For TRUB1 staining, TRUB1 Rabbit anti-human polyclonal, 50-172-8037, Protein tech; for PUS7 staining, PUS7 Rabbit anti-human, HPA024116, Sigma) in 1% BSA/PBS-Triton (0.1%) overnight at 4°C. The following day, the cells were washed with PBS-Tween 20 (0.1%) and incubated for one hour in 1:1000 secondary antibody (Mouse anti-rabbit IgG-PE-Cy7, NC1569698, Fisher) in 1% BSA/PBS-Triton (0.1%) at room temperature and stained using DAPI.

### siRNA Knockdown (KD) of TRUB1 and PUS7

SH-SY5Y cells were cultured in untreated media for 24h for KD and control samples. The media was replaced with siRNA delivery media (Horizon, B-005000-500) with 1uM of Accell Non-targeting Control Pool (Horizon, D-001910-10-05), PUS7 siRNA (Horizon, E-015341-00-0050) or TRUB1 siRNA (Horizon, E-016391-00-0050) in delivery media for KD samples and cultured for 72h. Total RNA was extracted after three days for qPCR KD confirmation and cells were fixed for IF imaging. The KD and scrambled control cells were stained with anti-Pus7 and anti-Trub1 antibodies to evaluate protein expression following the knockdown.

### Total RNA extraction

Total RNA was extracted from cells by combining a TRIzol (Invitrogen,15596026) RNA extraction and the PureLink RNA Mini Kit (Invitrogen, 12183025). Cells were first washed with ice-cold PBS, followed by incubation for 5 min in TRIzol at RT (2ml for 10mm dishes and 300ul for 8-well plates). Then, the solution was transferred to Eppendorf tubes, and 200ul chloroform (Thermo Scientific Chemicals, AC423555000) was added to each 1ml of TRIzol. The solution was mixed by shaking it for 15 sec and incubated at RT for 3 min, followed by centrifugation for 15 min at 12000 x g at 4°C. The aqueous supernatant was then transferred to a new tube, and the manufacturer’s recommended protocol was followed for PureLink RNA Mini Kit RNA extraction. RNA was quantified using the RNA Qubit assay.

### Poly-A RNA isolation

According to the manufacturer’s protocol, poly-A mRNA was selected using the NEBNext Poly(A) mRNA Magnetic Isolation Module (E7490L). RNA was quantified using the RNA Qubit assay.

### RT-qPCR

The extracted total RNA was treated with TURBO DNase (Fisher, AM2238) following the manufacturer’s protocol. The RNA is then reverse transcribed using SuperScript III RT kit (Fisher,18080044) using the target primers and housekeeping gene HPRT. qPCR was performed using Luna qPCR master mix (NEB, M3004).

### CMC-RT and ligation assisted PCR analysis of psi modification (CLAP)

The method is adapted from the protocol of Zhang et al. Primer3Plus was used to design primer sequences, which were tested for unspecific binding with BLAST.

### Direct RNA library preparation and sequencing

A direct RNA sequencing library (SQK-RNA002) was prepared following the manufacturer’s instructions. Briefly, 500ng poly-A tailed RNA was ligated to ONT RT adaptor (RTA) using T4 DNA ligase (NEB, M0202M) and reverse transcribed by SuperScript III Reverse transcriptase (Invitrogen, 18080044). The product was then purified using Agencourt RNAClean XP beads (Beckman, A63987) ligated to the RNA adaptor (RMX) and purified by Agencourt RNAClean XP beads, followed by washing with wash buffer (WSB) and eluted in elution buffer (ELB). The final product was mixed with an RNA running buffer and loaded into R9.4.1 FLO-MIN106D flow cells from ONT. For the KD samples and scrambled control, the samples were loaded onto PromethION flow cells (ONT, FLO-PRO004RA).

### Base-calling and alignment

Fast5 files were basecalled using Guppy version 6.4.2 and aligned to the human reference genome (hg38) using Minimap2 version 2.17 with the ‘‘-ax splice -uf -k 14’’ option. The aligned sam files were converted to .bam and indexed using samtools version 2.8.13.

### Paired IVT enrichment and psi detection with Mod-*p* ID

We generated an unmodified IVT transcriptome paired to the untreated library which we used as a control to identify putative psi positions for the. All the positions with a high number of reads in the DRS sample and insufficient coverage in the paired IVT library were enriched using a pan human IVT which merges IVT libraries from 6 human cell lines^68^. We apply a conditional approach to determine the baseline: we use the paired IVT when the number of reads is sufficient (>10), and we recover sites with insufficient reads in the paired IVT by using the pan-human IVT as a baseline. After defining the appropriate IVT control for each site, we filtered the putative psi positions detected by the Mod-p ID tool according to multiple criteria. First, we defined a psi site to be significant if at least two biological replicates had a p-value < 0.001. We then filtered all the significant sites for the number of reads in the DRS and IVT samples, to retain positions with at least 10 reads in both libraries. To account for the presence of SNVs, we filtered the significant sites with sufficient coverage to those with a U-to-C basecalling error < 10 in the IVT library.

### SNR Calculation

We modeled the IVT and DRS data separately for each target position with a beta-binomial distribution using Jeffrey’s noninformative prior^65–67^. We used U and C counts to parameterize the beta of the beta-binomial distribution and calculate the log marginal likelihood of the posterior distribution. Additionally, we modeled a combined distribution of IVT and DRS U and C counts with the same beta-binomial and Jeffrey’s prior. The ratio of these log marginal likelihoods approximates the degree to which the U to C mismatch at a position is better modeled with two independent distributions instead of a single joined distribution. This calculation is important because in DRS the total amount of reads is limited by the flow-cell design, therefore some abundant transcripts have very high coverage while others do not. For a transcript that is very highly abundant (i.e., >100 reads), a lower mismatch error could be considered significantly modified (i.e., <20% mismatch error). For a transcript that is lower abundance (∼10 reads), these sites could also be considered significant if most of the reads have a high mismatch error. According to Kass and Raftery’s model ^68^, we selected a value of SNR ≥ 1 to guarantee a strong DRS signal level as compared to the IVT.

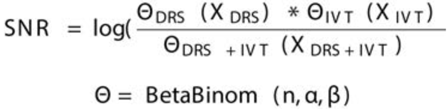

### SNR analysis and Mod-p ID cutoff selection in KD and perturbated libraries account for depth limitations

A MinION flow cell typically generates 1–2 million aligned reads^69^. This allows to achieve very high coverage depths of more than 50x in smaller genomes and transcriptomes (i.e. yeast and bacteria) but limits the coverage for the human transcriptome. Opposite to Illumina RNA-seq, the depth of a single DRS replicate cannot be improved with longer runs because of how the flow cell is constructed. When using DRS, we cannot increase the coverage/sequence depth by combining the sequencing output from different flow cells as each technological replicate is loaded on a unique flow cell, therefore merging the data coming from different flow cells cannot solve depth limitations. To leverage these technical depth limitations and guarantee a highly conservative and rigorous selection of psi sites, we propose an SNR-based filtration of the data to be coupled with the previously developed Mod-p ID tool. Therefore, we relax the U- to-C cutoff originally used for Mod-p ID as this is now coupled with the assessment of each site noise ratio, calculated using the unmodified IVT transcript. We apply more stringent cutoffs on the KD libraries as the selection of psi sites will affect the downstream analysis. The differential U-to-C mismatch calculated on KDs vs Scrambled library is particularly conservative as it requires a decrease of at least We then relax the U-to-C mismatch cutoff considering the CI of the perturbated libraries. We further apply a minimum read cutoff (>10) for each putative site in both the DRS and IVT sample. This criterion accounts for differences in depth between the smaller MinION libraries used in the original version of Mod-p ID, and the current PromethION runs. The read cutoff adds up to the original p-value filtration (p-value<0.001), to account for the increased number of reads per site in PromethION libraries. Although the selection of the cutoffs used in this work is determined empirically, we apply multiple filtering steps to account for coverage, SNVs presence, occupancy levels. We further validate the filtered list of sites with genetic and orthogonal methods. All these steps allow us to confidently define a rigorous, although small, list of psi-sites.

### Assignment of psi-sites to TRUB1 and PUS7 enzymes

We define sites as knocked down if they met the SNR ≥ 1 criteria, had sufficient coverage in KD and SCR libraries (>10) and a specified % decrease between U-to-C error in the control and KD sample. For positions with more reads (>30 reads), we set the cutoff at a 15% dicrease in occupancy in the KD-sample. For positions with <30 reads, we set the cutoff at a 30% decrease in occupancy. These cutoffs allow us to call a site as knocked down only when a substantial decrease in occupancy is observed the KD library vs the control but are selected to be more stringent with sites that have fewer reads.

### ML training on synthetic standards

We trained a ML model according to Makhamreh et al.^29^ for 10 synthetic RNA modified standards, of which four (PSMB2,PRPSAP1,MRPS14,MCM5) come from the original paper.

### Synthetic sequence design

We constructed six synthetic RNA oligos according to Tavakoli et al.^25^ and Gamper et al.^27,29,70^, each for a specific psi-site. Two versions of each RNA were prepared, one with 100% uridine and the other with 100% psi at the center of the synthetic standard.

## Resource availability Lead contact

Further information and requests for resources should be directed to and will be fulfilled by the lead contact, Sara H. Rouhanifard.

## Accessing Publicly Available Data Sets

All IVT Libraries used in this work were sourced from NIH NCBI SRA under BioProject accession PRJNA947135.

## Data and code availability

All fastq data for Direct Libraries generated in this work has been made publicly available in NIH NCBI SRA under the BioProject accession PRJNA1092333.

All bam files used for Mod-*p* ID can be found at the following Dropbox link: https://www.dropbox.com/scl/fo/4slozy1wmyxftbfzgyhaf/AE1nNQYNOoW-RUQibr0LTX8?rlkey=zqg24v8kpppst041d3oky0uk4&st=zu35f6i1&dl=0

All psi-sites detected by Mod-p ID in the KD libraries are available in **Supplementary Table 8**. The SNR valued calculated on KD libraries can be found in **Supplementary Table 9**. Knocked down psi-sites assigned to PUS enzymes are available in **Supplementary Table 10**.

All code can be found at github.com/RouhanifardLab/NeuronalEpitranscriptomePlasticity.

## Acknowledgments

S.H.R and M.W. acknowledge support from the National Institutes of Health (R01HG011087, R01HG012856). Y.M.H acknowledges support from the National Institutes of Health (R35-GM134931).

## Author contributions

Conceptualization O.F., S.T., S.H.R, and M.W.; Methodology O.F., S.H.R, and M.W.; Software O.F., S.T., A.M., and S.A.; Formal analysis O.F.; Investigation S.T., D.B., O.F., H.G., and Y.Q.; Resources S.T. and D.B.; Generation of Synthetic controls Y.M.H., H. G.; Writing - Original Draft S.T., O.S.F., S.H.R.; Writing - Review & Editing S.H.R., M.W., O.F., A.M., D.B., C.A.M., S.A, and K.Q.; Visualization O.F., S.T., S.H.R., K.N.; Supervision S.H.R., M.W., M.J. and Y.M.L.; Project administration S.H.R. and M.W.; Funding acquisition S.H.R.

