## Supporting Information for "Probing enzyme-dependent pseudouridylation using direct RNA sequencing to assess neuronal epitranscriptome plasticity"

*siRNA knockdown of TRUB1 and PUS7*

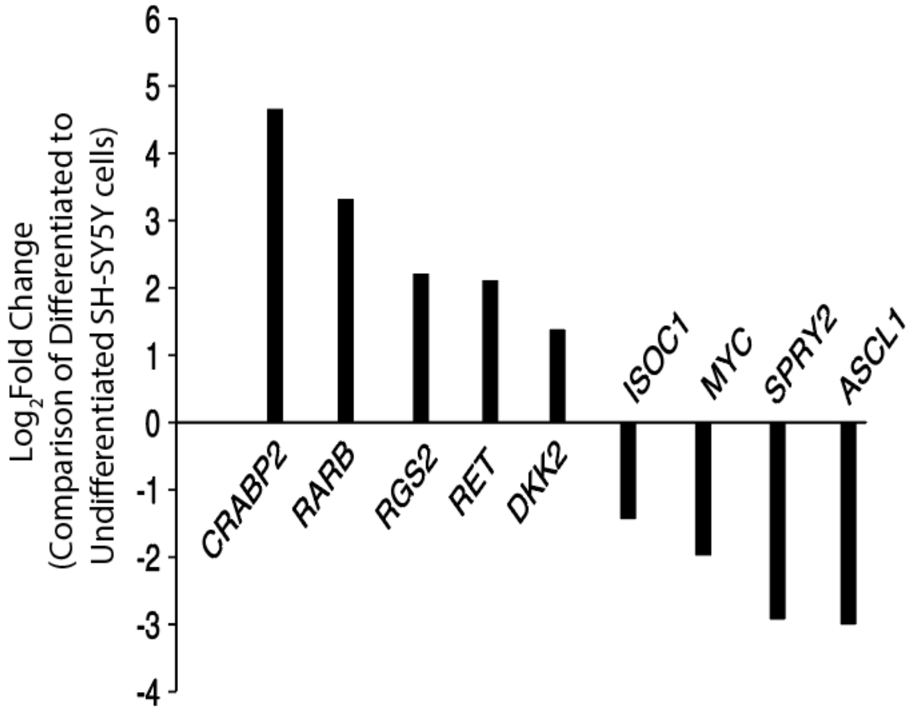

**Supplementary Figure 1.** Differential mRNA expression of known differentiation markers supports a change in the SH-SY5Y cellular state after the retinoic acid-induced differentiation treatment.

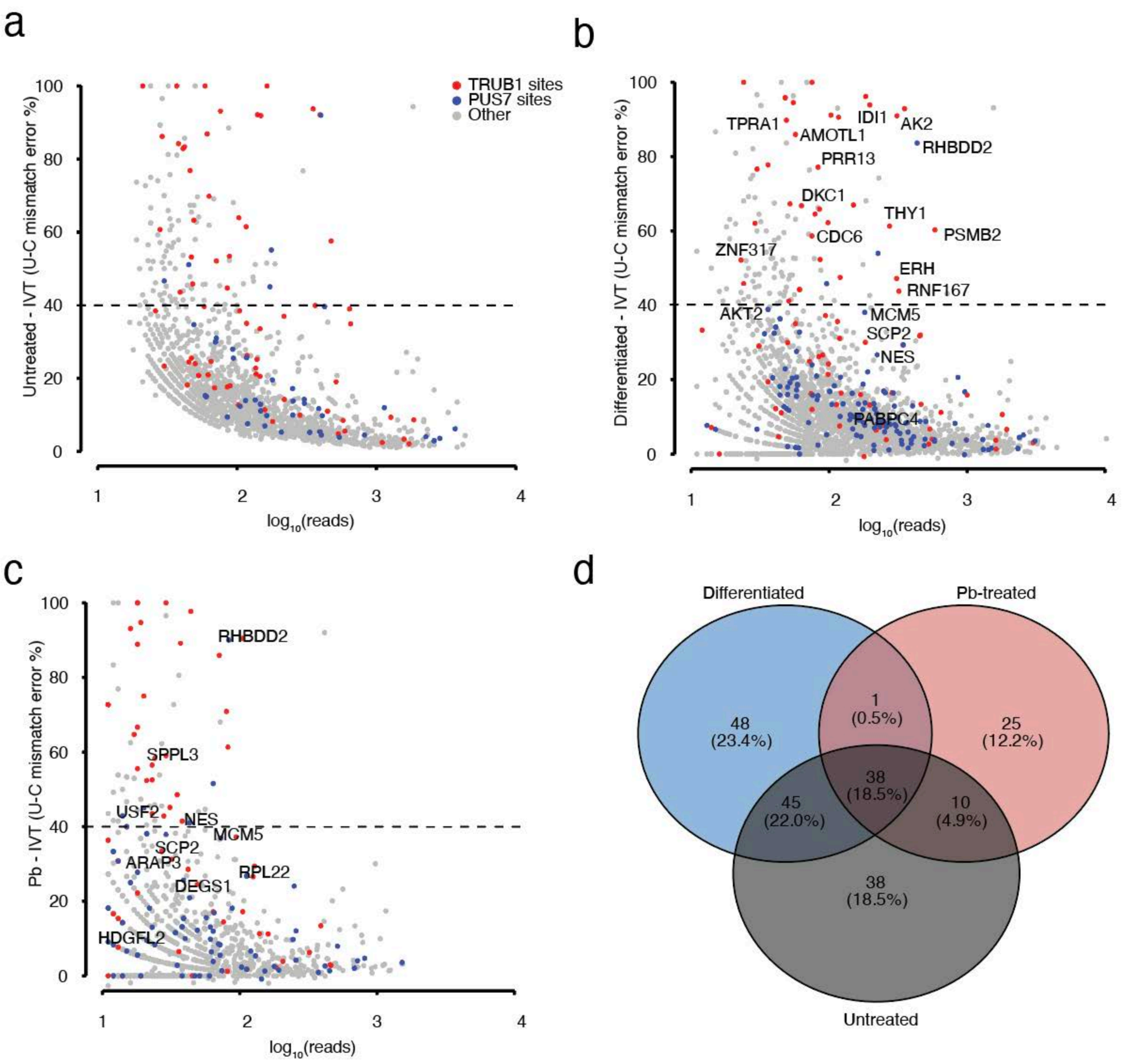

**Supplementary Figure 2**. Psi identification by Mod-p ID and hypermodified type I sites.
**a, b, c.** The U-to-C mismatches detected by nanopore sequencing for psi positions detected by Mod-p ID ($p < 0.001$) versus the log10(reads) of the unmodified, differentiated and lead-treated libraries. Targets with TRUB1 motif in red, targets with PUS7 motif in blue, any other motif in gray. Psi-sites are classified as hypermodified type I if their relative occupancy level is above 40% (dashed line).
**d.** Venn diagram on the hypermodified type I sites found in Undifferentiated, Differentiated and Lead-treated libraries shows the unique and common sites to the three groups.

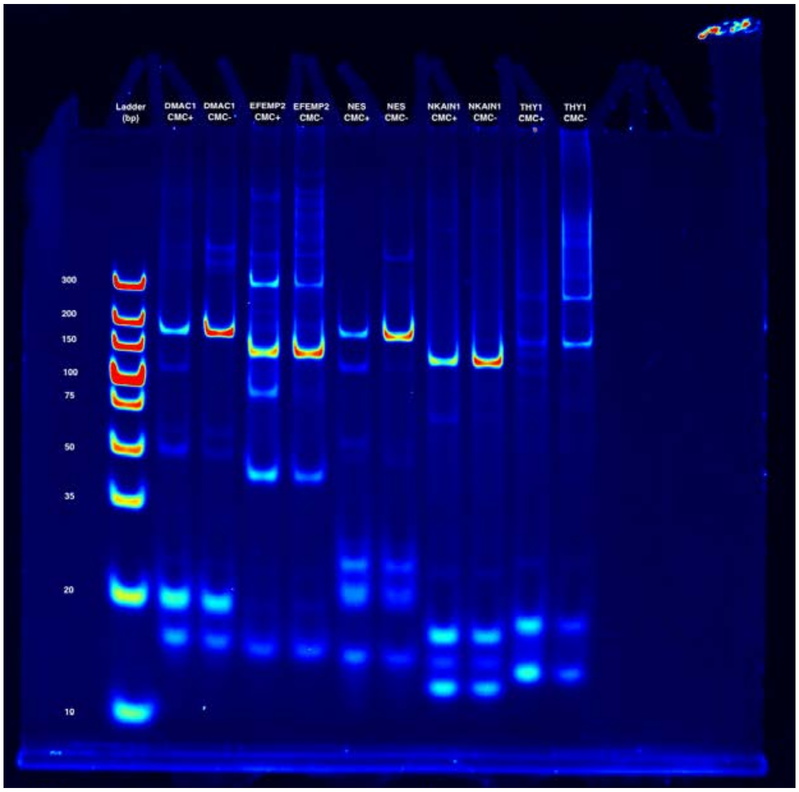

**Supplementary Figure 3.** CLAP validation of psi sites identified by mod-p ID in differentiated and/or lead treated SH-SY5Y cells that were only validated by the siRNA KD and not orthogonal method.

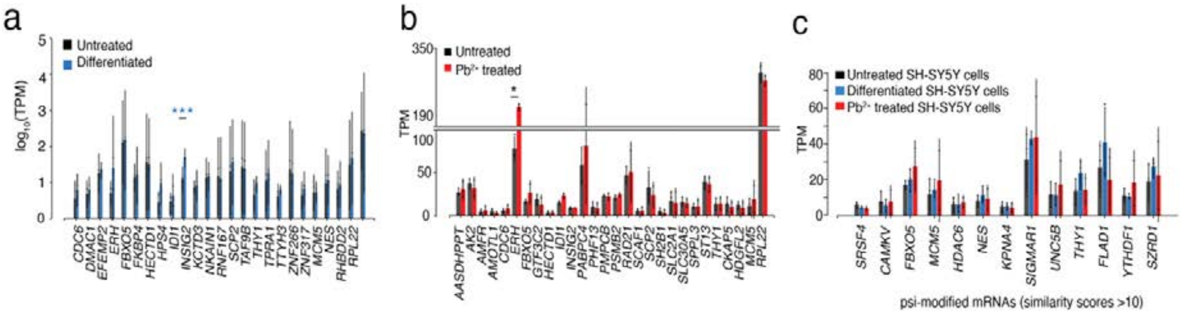

**Supplementary Figure 4.** mRNA expression levels for transcript bearing plastic psi-sites.
a. TPM of the transcripts bearing a validated psi modification that had the most significant differences between untreated and differentiated samples, determined by DRS.
b. TPM of the transcripts bearing a validated psi modification that had the most significant differences between untreated and lead-treated libraries.
c. TPM of the transcripts bearing a validated psi modification that had the most significant differences between all three conditions determined by DRS.
Individual colored bars represent each experimental condition, with error bars describing the standard error of the mean (SEM) across down sampled replicates. Individual replicates are shown as black dots. Statistics are performed by Student's t-test, comparing each KD group to the scrambled control sample. * $p < 0.05$, ** $p < 0.01$, *** $p < 0.001$.

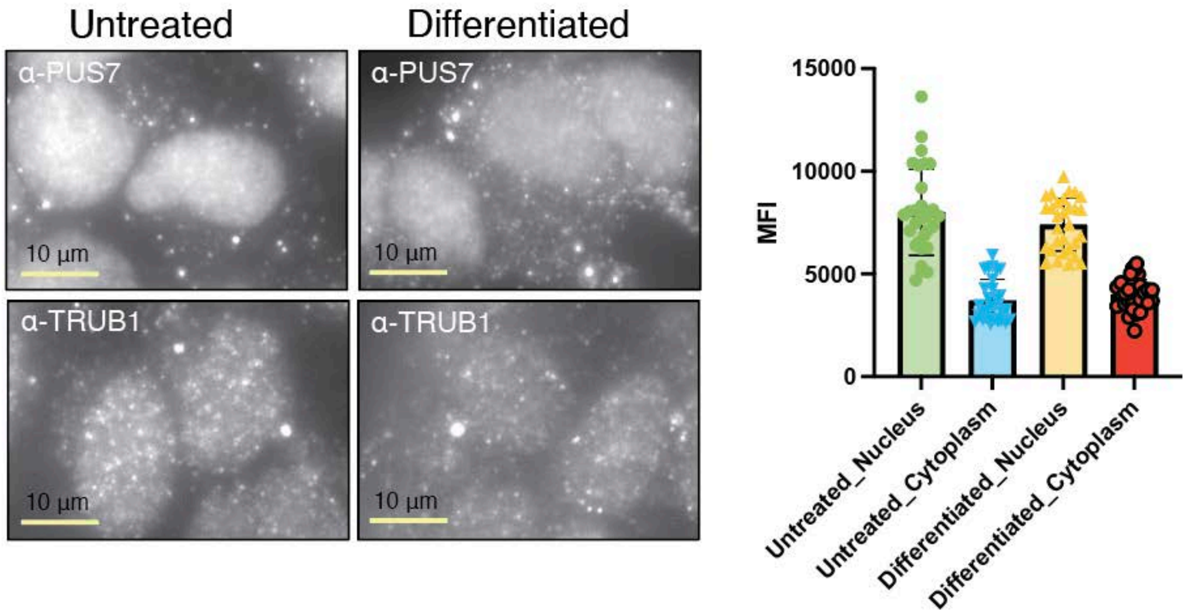

**Supplementary Figure 5.** IF images of untreated and differentiated SH-SY5Y cells using a. PUS7 and b. TRUB1 antibodies. (right) Comparison of PUS7 and TRUB1 antibodies' fluorescence intensity in the nucleus and cytoplasm of untreated and differentiated SH-SY5Y cells.

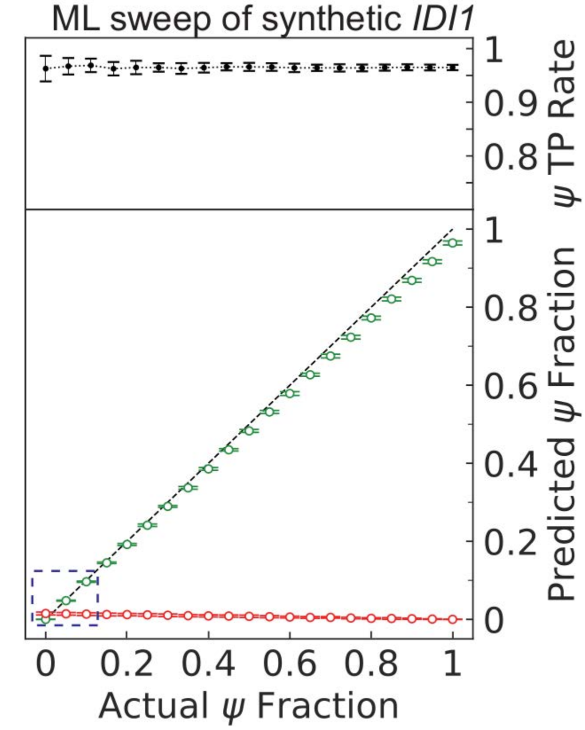

**Supplementary Figure 6.** ML classification results of model trained for I*DI1* with *IDI1* synthetic controls.
(Top panel) Mean±st. dev. of the true positive (TP) rate, also known as recall, for ψ, calculated as the number of TP calls divided by the sum of TP and false negatives (FN), as a function of the ψ fraction in the test set using 25 unique gradient boosting classification (GBC) models generated with repeated stratified 5-fold cross validation.
(Bottom panel) Mean ± st. dev. of the TP (green) and false-positive (FP, red) predictions for ψ, divided by the total number of samples in the test set as a function of ψ fraction for the same set of GBC models used in the top panel. The region where the frequency of FP calls remains consistently lower than the TP calls is highlighted in blue (approximately 5% ψ fraction).

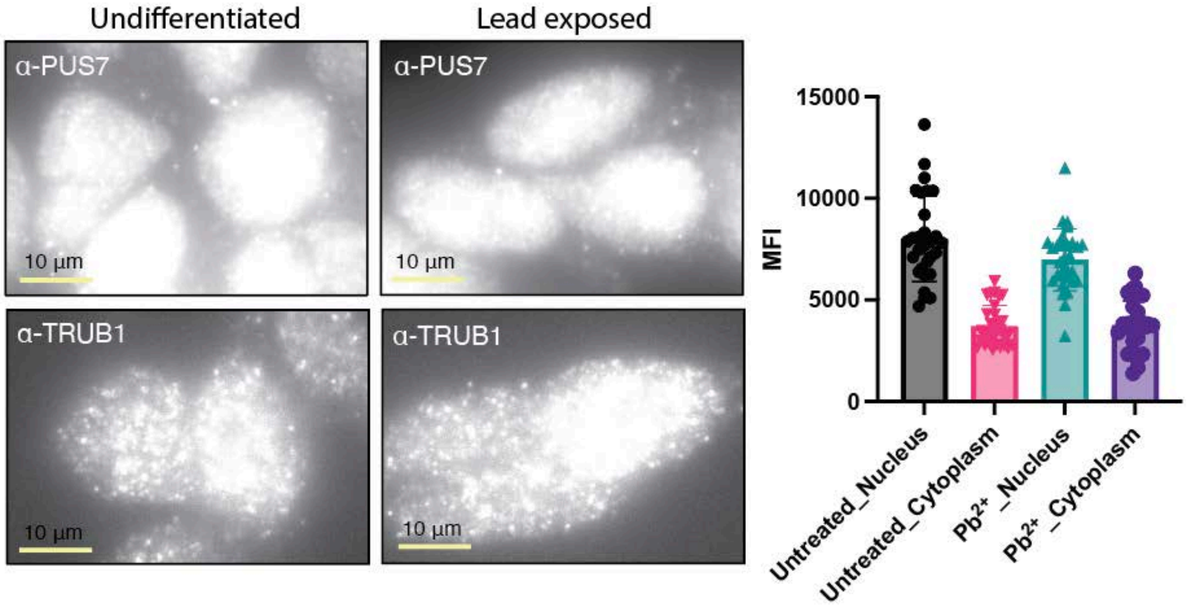

**Supplementary Figure 7.** IF images of untreated and lead-exposed SH-SY5Y cells using PUS7 and TRUB1 antibodies. (right) Comparison of PUS7 and TRUB1 antibodies' fluorescence intensity in the nucleus and cytoplasm of untreated and lead treated SH-SY5Y cells.

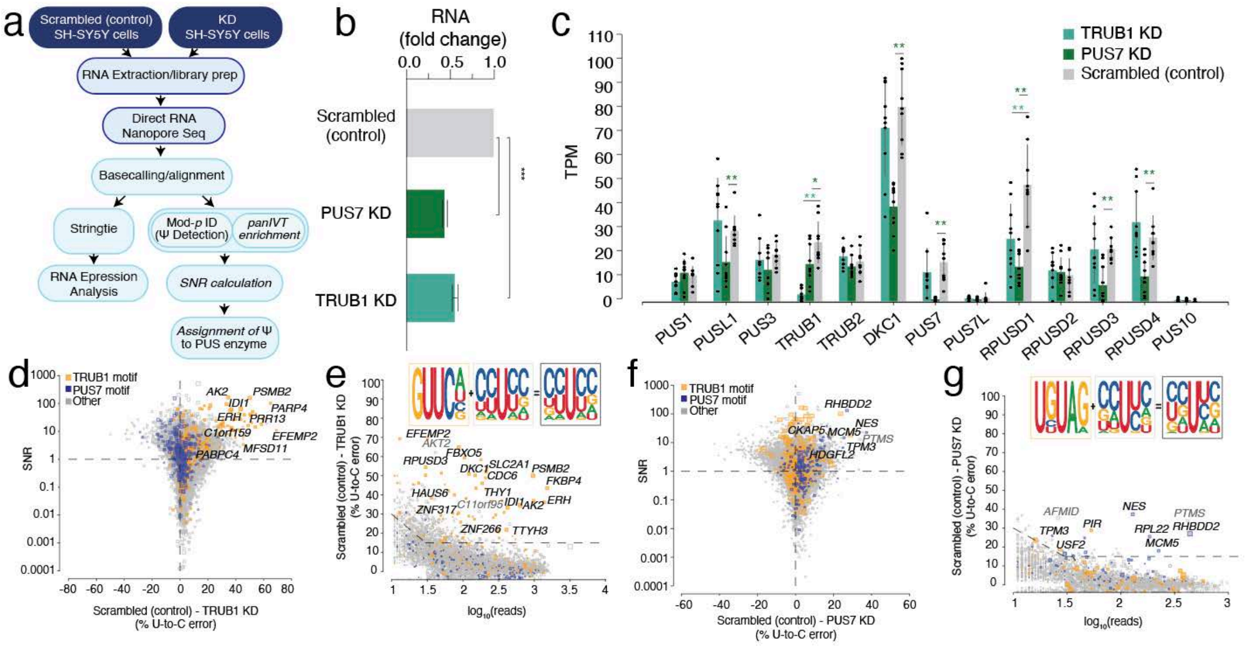

**Supplementary Figure 8.** Enzymatic KD of PUS enzymes and DRS is used to determine enzyme-mediated psi-sites.

a. Schematic workflow of siRNA knockdown (KD), DRS, and analysis.

b. The concentration of PUS7 and TRUB1 mRNA in SH-SY5Y cells for the scrambled (control), PUS7 KD, and TRUB1 KD, respectively, following siRNA KD, was quantified by RT-qPCR.

c. TPM of various PUS enzymes following TRUB1 knockdown (KD), PUS7 knockdown (KD), and scrambled (control) determined by DRS. Individual colored bars represent each experimental condition, with error bars describing the standard error of the mean (SEM) across downsampled replicates. Individual replicates are shown as black dots. Statistics are performed by Student's t-test, comparing each KD group to the scrambled control sample. * $p < 0.05$, ** $p < 0.01$, *** $p < 0.001$. P-values are shown

d. Signal-to-noise ratio (SNR) vs. the difference in U-to-C error % between Scrambled (control) and TRUB1 KD. Orange dots represent uridine positions within a TRUB1 motif, and blue dots represent uridine positions within a PUS7 motif.

e. Putative psi-positions determined by Mod-p ID are plotted according to the difference in U-to-C basecalling error in the scrambled (control) and the TRUB1 KD against the reads for each position. Inset shows the sequencing logo for positions within the TRUB1 motif, grey points above the threshold line, and total points above the threshold line.

f. Signal-to-noise ratio (SNR) vs. the difference in U-to-C error % between Scrambled (control) and PUS7 KD. Orange dots represent uridine positions within a TRUB1 motif, and blue dots represent uridine positions within a PUS7 motif.

g. Putative psi-positions determined by Mod-p ID are plotted according to the difference in U-to-C basecalling error in the scrambled (control) and the TRUB1 KD against the reads for each position. Inset shows the sequencing logo for positions within the PUS7 motif, grey points above the threshold line, and total points above the threshold line.

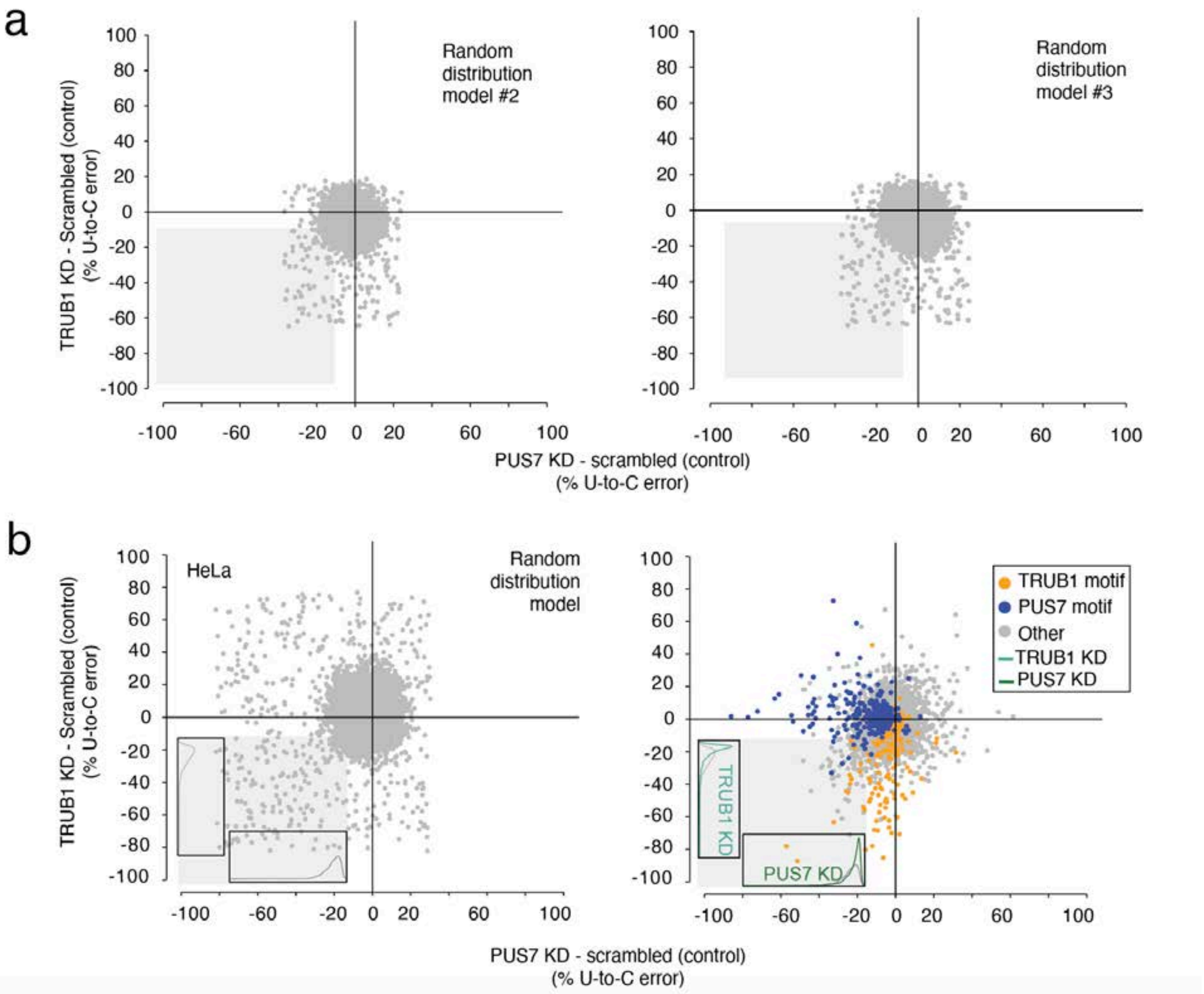

**Supplementary Fig.9** Co-regulation of TRUB1 and PUS7 enzymes.
**a.** Two additional random models of points constrained by the data parameters are used to test statistical significance with the real data on SH-SY5Y cell line. Statistics are performed on the III quadrant and performed with Mann–Whitney–Wilcoxon test.
**b.** Co-regulation effect of TRUB1 and PUS7 KD on HeLa cells. (Left) Random distribution model of points constrained by the data parameters. Distributions are calculated for points in the lower left quadrant as a function of each simulated knockdown. (Right) distribution of TRUB1 KD plotted against PUS7 knockdown. Distributions are calculated for points in the lower left quadrant as a function of each knockdown.

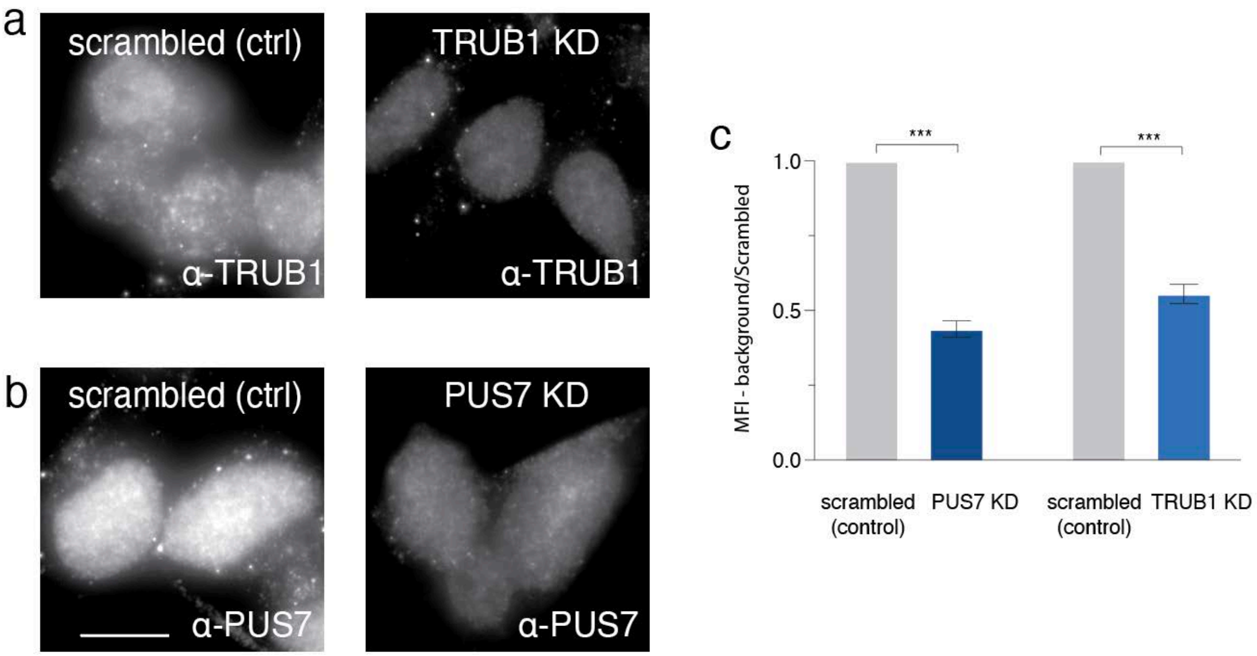

**Supplementary Figure 10.** Representative photomicrographs of a. TRUB1 knockdown (KD) and b. PUS7 KD vs Scrambled (control) cells show a substantial decrease in fluorescence in the KD library.

c. Immunofluorescence (IF) of TRUB1 KD, PUS7 knockdown KD, and siRNA control show a decreased mean fluorescence intensity (MFI) in the KD cells compared to the control. We observed a substantial fold-decrease in mean fluorescence intensity for Pus7 (0.56-fold ± 0.0275) as well as Trub1 (0.45-fold ± 0.0321) KD cells as compared to the scrambled siRNA control.
